# Identifying, phasing, and structurally annotating sex chromosomes for genome assemblies using CBS-tools

**DOI:** 10.64898/2026.09.23.753885

**Authors:** Lauren Whitt, Laramie M. Akozbek, Philip C. Bentz, Ellie E. Armstrong, Alex Harkess, Sarah B. Carey

## Abstract

A complete reference genome for species with chromosomally-determined separate sexes should contain scaffolds for all sex chromosome homologs. However, sex chromosomes present distinct computational challenges compared to autosomes. Here we present a *k*-mer based analysis that utilizes whole-genome sequencing of a few sex-identified isolates: Cytogenetics-By-Sequencing (CBS) tools. Unlike other approaches that typically address one aspect of the sex chromosomes, CBS-tools strives to guide users from the discovery of the heterogametic sex through identifying the sex-determination region (SDR). The core of CBS-tools is automated quantification of sex-specific *k*-mers in order to predict the heterogametic sex. Using publicly-available datasets, CBS-tools correctly identified the known sex-system of the 31 species tested. Additionally, we used these *k*-mers to verify and correct phasing of sex chromosomes between haplotypes in species representing different sex-systems. Finally, we used these *k*-mers to delimit the SDR boundary using an interactive web platform. CBS-tools was developed with previously unexplored sex chromosome systems in mind, but is also suitable for well-examined sex chromosome pairs.

## Introduction

Sex chromosomes have long fascinated biologists because of their role in sex-determination and the production of sexually dimorphic traits, their display of sex-specific patterns of molecular and genomic evolution, and the capacity to better understand species evolutionary histories. Sex chromosomes were first identified due to their cytological heteromorphy. In mealworms (*Tenebrio molitor*), males were observed to inherit a smaller chromosome in a homologous pair ^1^, which today we recognize as the Y chromosome in an XY system. Since their initial discovery, other sex chromosome systems have been identified that differ based on their patterns of inheritance and research has shown that they are not always cytologically distinct from each other or from other chromosomes. Within ZW systems, for example, females are heterogametic rather than males, a pattern that was demonstrated cytologically in the ruby tiger moth (*Phragmatobia fuliginosa*; ^2^. The first species in plants to have their sex chromosomes identified was a bottle liverwort (*Sphaerocarpos donnellii*; ^3^. In bryophytes, and other species that are haploid when they develop their reproductive structures, both sexes inherit a sex-specific chromosome (U in females, V in males), forming what is known as a UV system ^4^. Despite evidence of having separate sexes, many underlying sex chromosomes have yet to be discovered, in part owing to a lack of cytological heteromorphy ^5,6^ and the difficulty in identifying and assembling them bioinformatically ^7,8^.

Across this array of sex chromosome systems, a typical observation is a lack of recombination on a portion of the sex-specific chromosome(s) and its homologous pair, that is subtended by one or two pseudoautosomal regions (PAR). The non-recombining region is sometimes referred to as the sex-determining region (SDR), as genes that initiate sex-specific development have often been identified within this region ^9–11^. However, that is not universally the case, and multiple studies have suggested that the X or Z chromosome may contain the sex determination locus ^12–14^. As such, a more appropriate term may relate to the sex-specificity of the region (e.g., the male-specific region of the Y or female-specific region of the W). For simplicity, as we will discuss several different sex chromosome systems that span an array of biological variation, we will use SDR to refer to the non-recombining region of a sex chromosome. Importantly, this region of non-recombination is critical to precisely identify in order to computationally analyze sex chromosomes.

To investigate sex chromosomes, a primary goal of a genome assembly is to have a contiguous representation of the sex chromosome pair. Advances in genome sequencing and assembly technologies have improved the capacity to build high-quality references ^15–18^. Sex chromosomes, however, remain difficult regions of genomes to examine ^19^. There are three major challenges with building sex chromosomes in genome assemblies. First, because many species have homomorphic sex chromosomes ^5,6^ that cannot be distinguished via cytological approaches, this makes it difficult to identify the heterogametic sex ^5,6^. Second, once the heterogametic sex is known, accurate assembly and identification of the sex chromosomes can be difficult due to their unique characteristics. Non-recombining regions are typically repeat rich ^20–23^, can be structurally variable (e.g., large to small hemizygous INDELs and translocations) ^24^, and can have hyper-diverse levels of sequence divergence ^25,26^ that may impact the phasing into respective haplotypes. Third, the size of the SDR can vary drastically across systems ^5,6,27^, and delineating the boundaries of the PAR can be difficult because not all SDRs have a clear structural variant, like an inversion, that encompasses the entire region ^28^.

Here we present Cytogenetics-By-Sequencing (CBS), a comprehensive collection of tools that utilize *k*-mer based analyses and whole-genome sequencing (WGS) data from sex-identified individuals to identify, phase, and structurally annotate sex chromosomes. CBS-tools is available on GitHub at https://github.com/HudsonAlpha/CBS-tools.

## Methods

### Logic behind the tool

The CBS pipeline identifies and utilizes *k*-mers that are specific to one sex to identify, assist in phasing, and structurally annotate sex chromosomes. The basis of this approach (using an XY system as an example) is that the organellar genomes, autosomes, and X chromosomes are found in both sexes, while the Y is the sex-specifc chromosome inherited only by males. The Y typically contains one or two PARs, which recombine relatively freely with the X ^29^. Thus, the only region in an XY system that is male-specifc is the SDR. At a *k*-mer level, what this entails is that *k*-mers found in males, but not females, should be derived from the SDR (Y-mers). This logic equally applies to ZW systems wherein female-specific *k*-mers (W-mers) are derived from the SDR of the W. In haploid systems, where both females and males inherit sex-specific chromosomes ^4^, both sexes then should contain sex-specific *k*-mers that are derived from the SDR of the U (U-mers) and V (V-mers) chromosomes.

### WGS datasets

We sought out plant and animal species with well-demonstrated sex chromosome pairs in XY, ZW, or UV sex systems (Table S1) and publicly available WGS datasets for female and male isolates to validate the CBS-tools *k*-mer based approach. Following this criteria, we used WGS datasets for *Amborella trichocarpa* ^30^, thorny asparagus (*Asparagus horridus* ^31^), Greenland wolf (*Canis lupus* ^32^), cannabis (*Cannabis sativa* ^33^), papaya (*Carica papaya* ^34^), fire moss (*Ceratodon purpureus* ^35^), horse (*Equus caballus* ^36^), domestic cat (*Felis catus* ^37^), fig (*Ficus carica* ^38^), Pacific beach strawberry (*Fragaria chiloensis* ssp*. pacifica* ^39^), wild strawberries (*Fragaria virginiana* ssp*. platypetala* ^39^, and *Fragaria virginiana ssp. virginiana* ^39^), chicken (*Gallus gallus* ^40^), threespine stickleback (*Gasterosteus aculeatus* ^41^), Antiochus longwing butterfly (*Heliconius antiochus* ^42^), hop (*Humulus lupulus* ^13^), Japanese hop (*Humulus scandens* ^43,44^), red bayberry (*Morella rubra* ^45^), white mulberry (*Morus alba* ^46^), mulberry (*Morus notabilis* ^47^), platypus (*Ornithorhynchus anatinus* ^48^), lion (*Panthera leo* ^49^), date palm (*Phoenix dactylifera* ^50^), white poplar (*Populus alba* ^51^), balsam poplar (*Populus balsamifera* ^52^), black cottonwood (*Populus trichocarpa* ^52^), willows (*Salix cardiophylla* ^53^, and *Salix dunnii* ^54^), peat moss (*Sphagnum divinum* ^55^), spinach (*Spinacia oleracea* ^56^), and brown bear (*Ursus arctos* ^57^) (Table S1, Table S2). Isolates were grouped into males or females according to the reported phenotype. Illumina WGS reads were trimmed with Trimmomatic v0.39 ^58^ using the parameters: LEADING:3 TRAILING:3 SLIDINGWINDOW:10:30 MINLEN:40.

### Sex-determining system identification

The first step for examining sex chromosomes in a genome reference is to identify which sex contains the heterogametic pair when it is unknown. The CBS framework is founded and centered on identifying sex-specific *k*-mers with the CBS-Discovery function. CBS-Discovery uses *k*-mer counting functions from Meryl ^59^ to generate sex-specific *k*-mer lists for each sex. In each species’ WGS dataset, *k*-mer count databases are created for each isolate, and combined *k*-mer count databases of all male isolates and all female isolates, respectively (Figure 1). All CBS-Discovery results shown here use the standard 21-mers ^60^, though users may choose any *k* compatible with Meryl. Next, *k*-mer count databases between isolates of the same sex are intersected to return only *k*-mers shared between isolates. Finally, we take the difference between the shared *k*-mer intersects from each sex and all *k*-mers from the combined count databases of the opposite sex to return *k*-mers specific to only one sex. In an XY system, this difference should yield far more male-specific *k*-mers (Y-mers) and in an ZW system there would be many more female-specific *k*-mers (W-mers).

**Figure 1.**
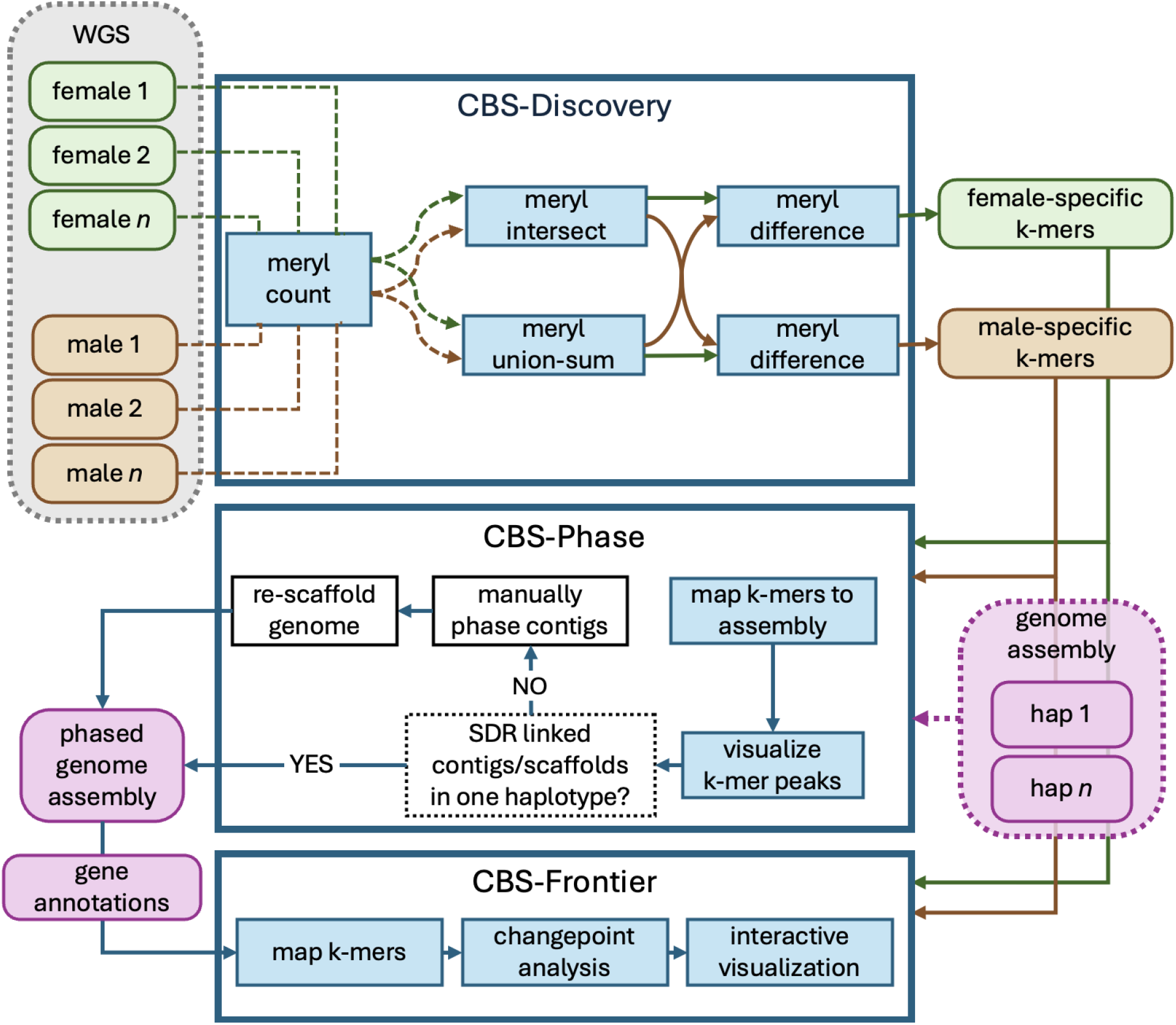
Overview of the CBS-tools functions, including options to fix sex-chromosome phasing across haplotypes and visualize SDR boundaries and gene models. Whole-genome sequencing from male and female isolates are put through CBS-Discovery’s *k*-mer based operations to identify male and/or female specific *k*-mers. Sex-specific *k*-mers can then be mapped to the genome assembly of a het-sexed individual to check that sex-linked contigs are properly phased in CBS-Phase. Steps performed automatically by CBS-tools are colored blue and steps that need to be performed manually by users are in white (no color). Finally, the properly phased genome assembly, sex-specific *k*-mers, and gene annotations (optional) can be imported into CBS-Frontier for interactive visualization of the sex-determining region and its features.

To ensure the resulting output of CBS-Discovery recovers *k*-mers from the SDR, we mapped the sex-specific *k*-mers to existing conspecific references of the heterogametic sex with BWA-MEM v0.7.19 using the parameters: -k 21 -T 21 -a -c 10 ^61^ (Table S1). The species for which we found suitable references to map the sex-specific *k*-mer lists were: Amborella ^30^, thorny asparagus ^31^, Greenland wolf ^62^, cannabis ^13^, papaya ^63^, fire moss ^35^, horse ^64^, domestic cat ^65,66^, fig ^67^, Pacific beach strawberry ^68^, chicken ^69^, threespine stickleback ^70^, Antiochus longwing butterfly (unpublished, GCA_964145975.1), hop ^13^, Japanese hop ^71^, red bayberry ^45^, white mulberry ^46^, mulberry ^47^, platypus ^18^, lion ^72^, date palm ^73^, white poplar ^74^, black cottonwood ^75^, willow ^54^, peat moss ^55^, spinach ^56^, and brown bear (Armstrong et al., *in review*). In a few cases a conspecific reference was not available; for balsam poplar, wild strawberries, and the willow *S. cardiophylla*, we instead used closely-related species (black cottonwood ^75^ Pacific beach strawberry ^68^, and *Salix arbutifolia* ^76^, respectively) with existing evidence of shared sex chromosome locations ^39,52,53^.

### Sequencing depth and isolate number subsets

We explored how WGS sequencing depth and the number of isolates used impacted our ability to identify and detect the sex system and SDR with CBS-tools. As reliably phased reference genomes of the heterogametic sex were not publicly available for all of the species tested in CBS-Discovery, we chose a representative subset with suitable reference genomes to explore depth and isolate number impacts in XY and ZW systems with different SDR sizes: Amborella ^30^, cannabis ^13^, chicken ^69^, threespine stickleback ^70^, hop ^13^, white mulberry ^77^, and brown bear (Armstrong et al., *in review*). We used Rasusa v2.2.2 ^78^ to randomly subset each isolate’s sequencing depth to 2.5✕ and 5✕. To test the number of isolates used, we chose the species with the largest pools of isolates in the XY and ZW systems for subsetting: threespine stickleback (XY) and Amborella (ZW). Subsets sizes of 3, 6, 12, and 24 males and females were randomly sampled 10 times from both species where possible. The only exception was the 24 isolate sampling in Amborella, where all 24 female isolates were used in all 10 replicate comparisons since there were only 24 total. To evaluate the effect of sequencing depth of coverage (x1) and number of isolates (x2) on the portion of *k*-mers mapping outside of the heterogametic-sex specific chromosome in all 31 species (y), we used beta regression models from the betareg R-package ^79^: (y ∼ x1 + x2) and estimated marginal means with the emmeans R-package ^80^.

### Phasing sex chromosomes in reference genomes

Genomic analyses of sex chromosomes rely on assembling fully-phased, contiguous genomic references. However, many current sequencing and assembly strategies do not reliably recover telomere-to-telomere chromosomes, nor can they accurately parentally-phase assemblies into respective haplotypes. As such, it can be unclear which contigs are sex-linked, which haplotypes they reside in, and whether they require manual curation prior to scaffolding. The CBS-tools suite includes a helper function, CBS-Phase, to identify SDR-linked contigs using the sex-specific *k*-mers from CBS-Discovery (Figure 1). The *k*-mer lists are mapped to each haplotype assembly of the reference genome using BWA-MEM v0.7.19 ^61^ and mapping rates are calculated with SAMtools v1.23.1 ^81^ and bedtools v2.31.1 ^82^. Additionally, users can opt to visualize average mapping depths across contigs in plots generated with ggplot2 ^83^. High *k*-mer mapping rates and covered bases indicate which contigs or scaffolds contain the SDR to aid users in the manual sorting of SDR-linked contigs across haplotypes assemblies. These methods were previously demonstrated by the authors in the hop ^13^, cannabis ^13^, and Amborella ^30^ genomes, and are now partially automated within CBS-Phase. In this paper, we present these three assemblies post phasing with the manual joint-scaffolding step (example described below) outlined in their respective citations.

The current gold-standard for phasing of a genome is through a trio-binning approach, where both parents’ and the offspring’s sequences are available. Trio-binning is able to phase maternal and paternal haplotypes ^84^, including the sex chromosomes. Although Hi-C enables chromosome-scale phasing in the absence of parental data, haplotype assignment is generally less accurate than with trio-binning because phase must be inferred indirectly from patterns of chromatin contact rather than directly from parent-specific sequence variants. Consequently, Hi-C-based assemblies may contain more errors or regions of uncertain phase, particularly in regions with low heterozygosity or with complex genomic architecture. However, because trio-binning might not be a viable approach for many genome assembly projects (e.g., wild-collected accessions), we aimed to test whether CBS-tools recovers the sex chromosomes using Hi-C data. Using the data for brown bear “Adak”, which had both parental WGS data and Omni-C data available to phase the assembly ^57,85^; Armstrong et al., *in review*), we mapped the sex-specific *k*-mers from CBS-Discovery to both the released trio-binned assembly and a genome assembly for “Adak” using the Omni-C data for phasing instead. For the Omni-C-phased assembly, we used the same version of HiFiasm ^84^ as in the released trio-binned assembly (v0.16.0), the only difference being that we invoked the Omni-C option for phasing rather than providing trio data. We then mapped the male-specific *k*-mers to both haplotypes of the Omni-C assembly with CBS-Phase.

In its current design, CBS-tools does not directly identify the homogametic sex chromosome, in this case, the X. Therefore we demonstrated, as an example, the joint-scaffolding approach that works within the Hi-C contact map to validate the contiguity of the Y-mer mapped contigs, identify the PARs, and identify the X-linked contigs. We concatenated the contigs for the two Omni-C phased haplotypes for “Adak” together and scaffolded them using YaHS v1.1 ^86^. Prior to running YaHS, we mapped the Omni-C data to the contigs using BWA-MEM v0.7.17 ^61^ using the flag -5SP, PCR duplicates were marked with SAMBLASTER v0.1.26 ^87^, and finally we used SAMtools view v1.10 ^81,88^ with parameters: -S -h -b -F 2326. We used Juicer Tools v1.9.9 to generate the Hi-C contact map, and manually curated the X/Y scaffolds using JuiceBox Assembly Tools ^89^, to identify the underlying X- and Y-linked contigs. Contigs identified as sex-linked in the Omni-C phased assembly were mapped to the trio-binned assembly using minimap2 v2.30 ^90^, and we subsequently identified the coverage using SAMtools coverage ^81,88^.

### Interactive visualization for structural annotation of sex chromosomes

A critical next step for analyzing sex chromosomes is delimiting the boundary of the non-recombining SDR from the PAR. CBS-tools includes another helper tool, CBS-Frontier, that uses sex-specific *k*-mers produced from CBS-Discovery to predict the SDR boundary and identify gene models overlapping *k*-mer alignments within the genome (Figure 1). Using the indexed alignment files and reference index files, which can be generated by CBS-Phase or other alignment methods, the preprocessing scripts of CBS-Frontier compute mean per-base *k*-mer coverage via bedtools coverage ^82^ in 1 kb, 10 kb, and 50 kb fixed windows, allowing for additional coverage stratification by mapping quality (MAPQ) and primary or secondary alignments per window if applicable. If gene and repeat General Feature Format files are available, the intersection of these features with mapped *k*-mers can be acquired during preprocessing, where the user can adjust the minimum *k*-mer count thresholds for identifying overlaps using the flag ‘--min-kmer-count’. Users can also convert non-standard annotation formats into BED6 format to load an additional track for visualization. Following preprocessing, the output files can be loaded to the CBS-Frontier application for visualization. This application was built with React (v19.2.0) and TypeScript (v5.9.3) on the Vite (v7.1.11) system and deployed via Vercel. All calculations and functions are computed on-demand, and no user-uploaded information leaves the browser. In this application, users are able to interactively navigate multiple views (Genome, Region, Chromosome and Compare) and export plots as well as in-window gene and repeat tables. Furthermore, CBS-Frontier can estimate the boundaries of their putative SDRs using binary-segmentation changepoint detection algorithms at the 1 kb resolution (‘ruptures’, L2 or Normal cost model) ^91^.

We used the reference genomes of the heterogametic sex from the same representative species used to evaluate sequencing depth impact to demonstrate the SDR identification capabilities of the CBS-Frontier. We also included two of the species from the Salicaceae that have undergone SDR translocations. For each species, we generated alignment inputs with CBS-Phase using their respective heterogametic sex-specific *k*-mer lists identified with CBS-Discovery. To demonstrate CBS-tools Frontier visualization capabilities, we focused on a subset of species with demonstrated sex-determining genes, the brown bear (*SRY* ^9,10^), three-spined stickleback (*amhy* ^11^), and two of the Salicaceae species (black cottonwood and *S. dunnii*; *ARR17* ^92^). As the *ARR17* genes in the SDR of black cottonwood and *S. dunnii* were missing from the available gene annotation files, we used the BLAST tool in Phytozome 14 ^93^ to identify their locations with the gene Potri.019G133600 as the query.

## Results and Discussion

### CBS-Discovery creates sex-specfic k-mer lists

Data were sourced from nine animal and 22 plant species with previously well-demonstrated sex chromosome pairs. These 31 species span the cytological variation of sex chromosomes, with representatives from all inheritance types that have a sex-specific chromosome, as well as some multiple systems (Table S1). These data also largely represent independent evolutions of sex chromosomes, especially within the plants, with a vast range of SDR sizes, providing a robust test for the CBS-tools pipeline. The core of CBS-tools is cross-referencing *k*-mer lists across a collection of male and female isolates to obtain sex-specific *k*-mers. *K*-mers from the SDR should be specific to the heterogametic sex and not found in the homogametic sex. The results of CBS-Discovery follow the expected patterns for the sex chromosome cytotype for all species examined (Figure 2). The species with observed XY cytotypes yielded more male-specific *k*-mers identified, and conversely ZWs resulted in more female-specific.

**Figure 2.**
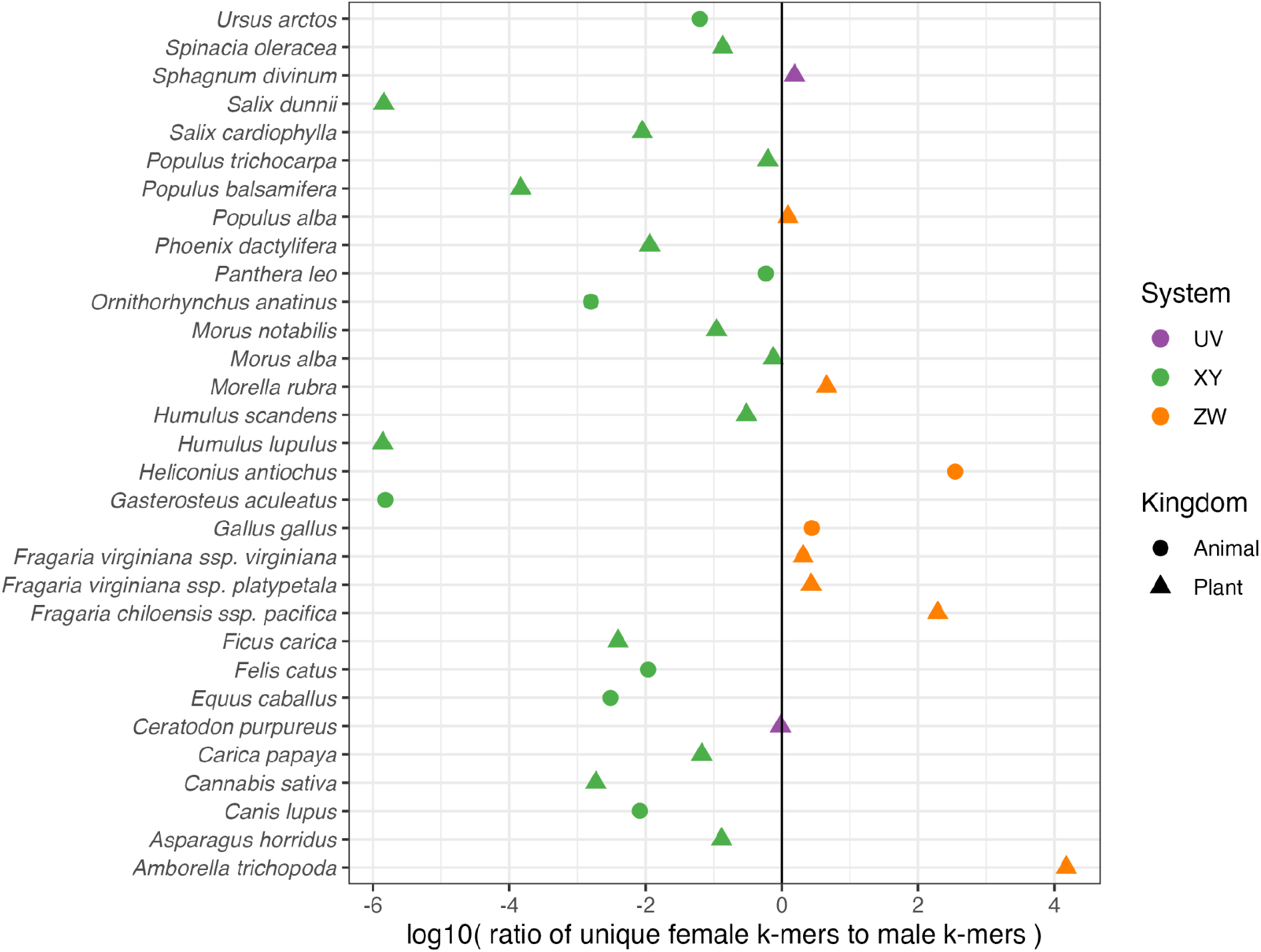
Log ratios of female:male unique *k*-mers from CBS-Discovery for various plant and animal species. Species are colored by their reported sex-system in primary literature, and species with many chromosomes (XYY, ZZW, etc.) have been summarized under XY or ZW (Table S1).

In this study, all sex-specific *k*-mer lists identified with CBS-Discovery aligned with the defined sex-specific chromosome (Table 1, Table S3, Figures S1-S2) regardless of sex-system, chromosome structure, number of sex-specific chromosomes (platypus X_1_X_2_X_3_X_4_X_5_Y_1_Y_2_Y_3_Y_4_Y_5_, Japanese hop X_1_Y_1_Y_2_, Antiochus longwing butterfly Z_1_Z_2_W_1_) or SDR size, with even small SDRs of 37kb to 140kb detected (Figure 3, Figures S1-S2). However, both biological and technical differences across systems may impact the recovery of sex-specific *k*-mers and the identification of each region. For example, the completeness of the genome reference is a technical constraint for further vetting the lists, as some existing references are not fully haplotype-phased nor completely and contiguously assembled (and some species lack a conspecific reference; see Methods, Table S1). We also note that mapping sex-specific *k*-mers to the genomes of different, but related, species is not ideal in many circumstances (Table S1), as *k*-mers between species can diverge, affecting mapping ability.

**Table 1.** Mapping rates of sex-specific *k*-mers to assemblies with decreasing sequencing genome sequencing depth inputs to CBS-Discovery. SDR boundaries estimated by changepoint analysis (Fig 3; STable 4). SDR boundaries for mapping as determined by changepoint analysis. Mapped reads are counted as primary mappings only. For assemblies split by haplotype, reads mapping to one set of autosomal chromosomes and both sex chromosomes are shown. Instances where CBS-Discovery returned 0 sex-specific *k*-mers changed to 1 for calculating the female to male (F:M) ratios.

| Species | F:M ratio | Female-specific |  |  | Male-specific |  |  |
| --- | --- | --- | --- | --- | --- | --- | --- |
| | | Total $k$-mers | Mapped | Within SDR | Total $k$-mers | Mapped | Within SDR |
| *A. trichopoda* |  |  |  |  |  |  |  |
| Full WGS | 14,937 | 14,937 | 99.80% | 99.80% | 0 | 0% | 0% |
| 5x normalized | 1,514 | 1,514 | 100.00% | 100.00% | 0 | 0.00% | 0.00% |
| 2.5x normalized | 24 | 24 | 100.00% | 100.00% | 0 | 0.00% | 0.00% |
| *C. sativa* |  |  |  |  |  |  |  |
| Full WGS | 0.002 | 6,470 | 5.60% | 0.00% | 3,458,566 | 94.73% | 94.43% |
| 5x normalized | 0.005 | 18,276 | 20.25% | 0.00% | 3,770,827 | 99.74% | 99.21% |
| 2.5x normalized | 0.032 | 20,203 | 67.22% | 0.00% | 625,696 | 99.47% | 97.26% |
| *G. gallus* |  |  |  |  |  |  |  |
| Full WGS | 3 | 21,399 | 53.80% | 42.23% | 7,816 | 74.01% | 0.00% |
| 5x normalized | 4 | 26,126 | 71.17% | 62.72% | 5,841 | 79.82% | 0.00% |
| 2.5x normalized | 6 | 17,814 | 74.17% | 66.72% | 3,104 | 89.40% | 0.00% |
| *G. aculeatus* |  |  |  |  |  |  |  |
| Full WGS | 0.000002 | 0 | 0% | 0% | 657,493 | 96% | 96% |
| 5x normalized | 0.000058 | 0 | 0.00% | 0.00% | 17,369 | 97.27% | 97.24% |
| 2.5x normalized | 0.000268 | 0 | 0.00% | 0.00% | 3,731 | 94.80% | 94.72% |
| *H. lupulus* |  |  |  |  |  |  |  |
| Full WGS | 0.000001 | 1 | 100.00% | 0.00% | 717,627 | 92.61% | 91.50% |
| 5x normalized | 0.000089 | 33 | 100.00% | 0.00% | 372,507 | 92.45% | 91.33% |
| 2.5x normalized | 0.010098 | 545 | 99.63% | 0.00% | 53,972 | 91.85% | 90% |
| *M. alba* |  |  |  |  |  |  |  |
| Full WGS | 0.750 | 751872 | 45.95% | 0.01% | 1002767 | 43.39% | 13.45% |
| 5x normalized | 0.626 | 619199 | 54.17% | 0.01% | 989756 | 45.29% | 13.84% |
| 2.5x normalized | 0.646 | 561414 | 61.71% | 0.04% | 868649 | 51.45% | 13.54% |
| *U. arctos* |  |  |  |  |  |  |  |
| Full WGS | 0.063 | 161316 | 0.07% | 0.00% | 2570696 | 79.35% | 73.53% |
| 5x normalized | 0.117 | 181070 | 17.84% | 0.00% | 1553206 | 96.68% | 84.93% |
| 2.5x normalized | 0.889 | 750716 | 83.86% | 0.00% | 844406 | 96.13% | 72.05% |

**Figure 3.**
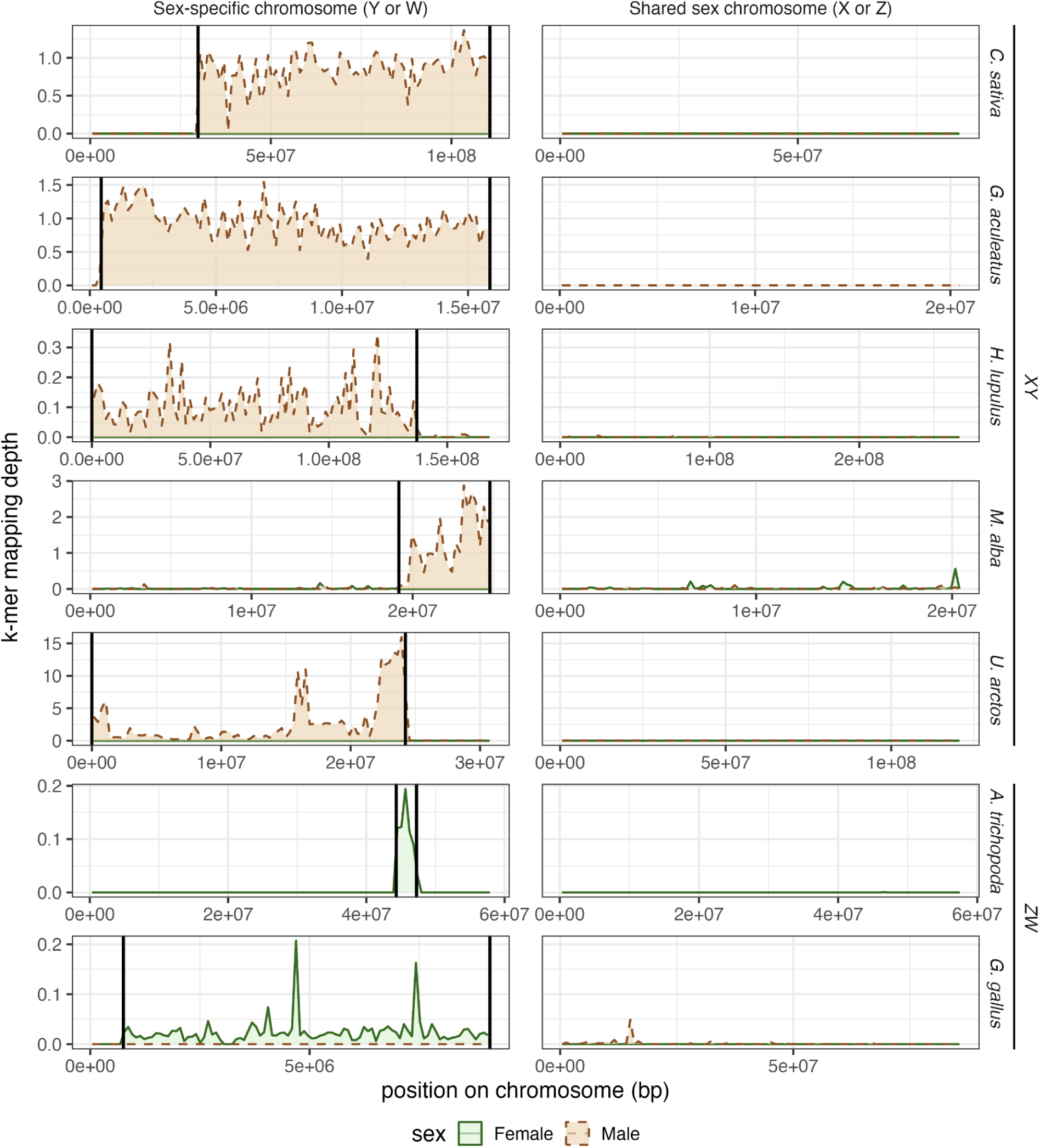
Mapping sex-specific *k*-mers to identify sex-determining regions (SDR) in representative subset: *Amborella trichopoda, Cannabis sativa* (cannabis), *Gallus gallus* (chicken), *Gasterosteus aculeatus* (threespine stickleback), *Humulus lupulus* (hop), *Morus alba* (white mulberry), and *Ursus arctos* (brown bear). Chromosomes are binned into 100 equal sized windows to calculate average *k*-mer mapping depths. Vertical black lines mark the estimated SDR boundaries in the heterogametic-specific chromosome as determined by changepoint analyses within CBS-Frontier (Table S5). For sex-specific *k*-mer mapping across all scaffolded chromosomes, see Figures S1-S2.

In our seven representative species where we more thoroughly explored the SDR, we found that the majority of heterogametic-sex *k*-mers map to the reported SDR, excluding white mulberry (Table 1). Moreover, CBS-Discovery identified *k*-mer lists that track the translocation of the SDR across the five Salicaceae species tested here (Table S1). In black cottonwood, balsam poplar, and the two willow species, we recovered more male-specific *k*-mers (Figure 2), which is consistent with the known XY chromosomes in these species. Importantly, the male-specific *k*-mers also hit the previously identified Y chromosome, with black cottonwood and balsam poplar both aligning to Chr19, *S. dunnii* aligning to Chr7, and *S. cardiophylla* aligning to Chr15 (Figure S1). In white poplar we recovered more female-specific *k*-mers, consistent with the evidence for a ZW system, and the *k*-mers aligned to Chr19 as expected.

We evaluated the effects of two technical variables that researchers have moderate control over when using CBS-Discovery: the number of isolates and sequencing depth of coverage in the datasets. The number of isolates used had the greatest impact (b = -0.034, z = -3.0, p = 0.003), with datasets having fewer isolates tending to show greater noise of *k*-mers mapping outside of the sex chromosome (Table 2, Table S4). This is likely due to the smaller amount of isolates not fully capturing population structure and unsampled diversity within the species, and thus returning rare or sub-population specific autosomal variation as sex-linked, because the homogametic sex does not possess these *k*-mers to filter them out of the heterogametic sex-specific *k*-mer list. Indeed, when using random samples of 3 isolates per sex in CBS-Discovery, roughly 21% of mapped female-specific *k*-mers are outside of the W-SDR in Amborella compared to less than 2% of female-specific *k*-mers when using 6 or more isolates per sex. Though the difference is less extreme in threespine stickleback, we also observe 7% of mapped male-specific *k*-mers mapping outside the Y-SDR when using 3 isolates compared to less than 1.5% when using 6 or more isolates (Table 2). Additionally, the random subsets of 3 isolates per sex return orders of magnitude more homogametic-sex *k*-mers than when using 6 isolates per sex in both species (Table 2). This trend holds as the number of isolates increases up to 24 isolates where no *k*-mers were uniquely found in the homogametic sex (Table 2). The CBS methodology is designed around a sampling of multiple male and female isolates in an attempt to capture as much autosomal genetic variation as possible in order to prevent their inclusion in putative sex-specific *k*-mer lists. There is no universal threshold of isolate needed to detect the SDR currently, as this threshold will be influenced by the species’ unique genetic diversity and evolutionary history. For species with conserved sex chromosomes, such as Amborella and threespine stickleback, more isolates will typically improve the signal, where the sex-specific *k*-mer ratios move away from 1.0 as more isolates are used in CBS-Discovery (Figure 4; Table 2).

**Figure 4.**
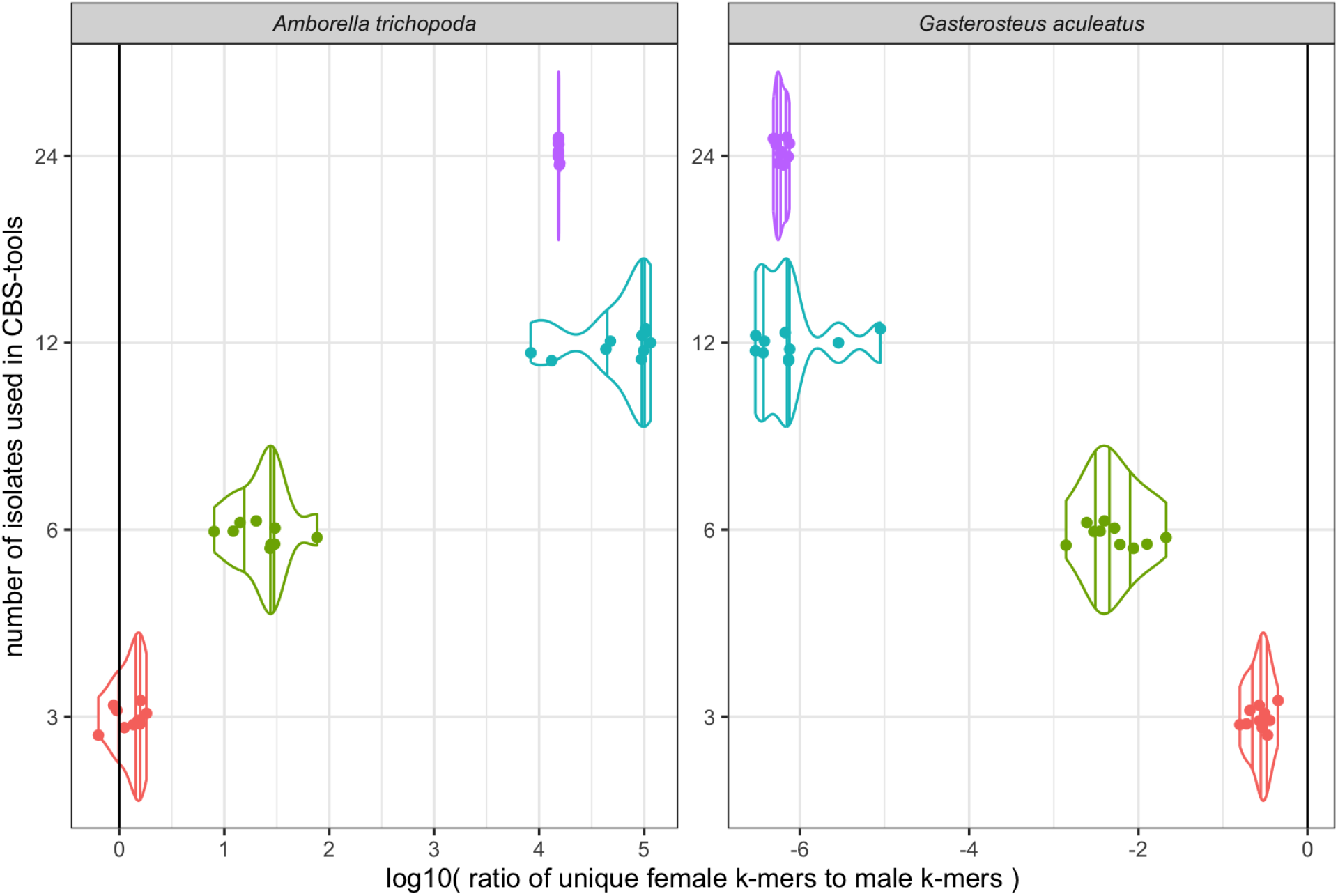
Comparing distributions of male:female unique *k*-mer ratios when using different numbers of isolates as inputs for CBS-Discovery. Random samples of 3, 6, 12, and 24 isolates were taken from both sexes in *Amborella trichopoda* (ZW system) and *Gasterosteus aculeatus* (threespine stickleback, XY system), except for the 24 isolate subsampling, where the same 24 *A. trichopoda* females were used since there were only 24. Ratios displayed in log10 scale.

**Table 2.** Average mapping rates to assemblies and the SDR when sex-specific *k-*mers are called with different numbers of male (M) and female (F) isolates. SDR boundaries estimated by changepoint analysis (STable 4). Mapped reads are counted as primary mappings only. For assemblies split by haplotype, reads mapping to one set of autosomal chromosomes and both sex chromosomes are shown. Instances where CBS-Discovery returned 0 sex-specific *k*-mers changed to 1 for calculating the female to male (F:M) ratios. Random samples n = 10, values displayed as mean ± std.err.

| Species | F:M ratio | Female-specific |  |  | Male-specific |  |  |
| --- | --- | --- | --- | --- | --- | --- | --- |
|  |  | Total *k*-mers | Total Mapped | Within SDR | Total *k*-mers | Total Mapped | Within SDR |
| *A. trichopoda* |  |  |  |  |  |  |  |
| 3M vs 3F | 1.16 | 1.7E+06 ± 9.5E+04 | 49.0% ± 4.8% | 27.6% ± 0.6% | 1.5E+06 ± 2.6E+05 | 28.1% ± 3.6% | 0.0% ± 0.0% |
| 6M vs 6F | 19.90 | 3.0E+05 ± 1.1E+04 | 96.5% ± 2.9% | 94.6% ± 2.7% | 1.5E+04 ± 2.7E+03 | 26.3% ± 5.7% | 0.0% ± 0.0% |
| 12M vs 12F | 23757.72 | 9.9E+04 ± 2.6E+03 | 99.9% ± 2.6% | 99.7% ± 2.6% | 4.2E+00 ± 1.7E+00 | 44.0% ± 44.0% | 0.0% ± 0.0% |
| 24M vs 24F | 15416.30 | 1.5E+04 ± 4.7E+01 | 99.7% ± 0.3% | 99.6% ± 0.3% | 0 | NA | NA |
| *G. aculeatus* |  |  |  |  |  |  |  |
| 3M vs 3F | 0.28 | 1.8E+06 ± 2.3E+05 | 29.5% ± 3.9% | 0.0% ± 0.0% | 6.3E+06 ± 3.2E+05 | 75.4% ± 2.8% | 69.0% ± 1.9% |
| 6M vs 6F | 0.01 | 2.8E+04 ± 8.7E+03 | 26.3% ± 5.9% | 0.0% ± 0.0% | 3.9E+06 ± 1.3E+05 | 95.4% ± 3.2% | 95.1% ± 3.1% |
| 12M vs 12F | 0.000002 | 6.1E+00 ± 3.3E+00 | 51.2% ± 45.8% | 0.0% ± 0.0% | 2.9E+06 ± 8.9E+04 | 95.9% ± 2.9% | 95.8% ± 2.9% |
| 24M vs 24F | 0.000001 | 0 | NA | NA | 1.7E+06 ± 8.1E+04 | 96.0% ± 4.6% | 96.0% ± 4.6% |

The depth of coverage of the isolates’ sequencing appears to have a lesser effect on *k*-mer noise (b = -0.017, z = -2.073, p = 0.038), with CBS-Discovery still returning the expected results in these lower depth datasets (Table 1), as the percentage of heterogametic specific *k*-mers mapping to the estimated SDR boundaries remains relatively unchanged across the seven example species. However, dropping to 5✕ and 2.5✕ sequencing depth does introduce more noise that can confound sex-chromosome identification. Naturally, low depth WGS datasets will reduce the likelihood of all unique *k*-mers being represented in an isolate’s sequencing due to the random chance of some portions of the genome not being represented. In CBS-Discovery, this can 1) reduce sex-linked *k*-mer signal due to some isolates missing sex-linked *k*-mers that are truly shared with the other same-sexed isolates, and 2) increase background noise (i.e. *k*-mer mapping peaks outside of the SDR) due to one sex missing universally shared *k*-mers from autosomes or the PAR that will then be misidentified as specific to the other sex. This can result in the sex-specific *k*-mers ratio approaching closer to one (Figure 2, Table 1), making it difficult to confidently identify the sex-system. We can see how these two problems play out when mapping *k*-mers from low depth sequencing after CBS-Phase. 1) Fewer heterogametic sex-specific *k*-mers are identified (Table 1). 2) Excluding chicken, we find a slight increase of heterogametic sex-specific *k*-mers mapping outside of the SDR, as defined by CBS-Frontier (Table S5) in the 2.5✕ sequencing depth of coverage datasets (Table 1); however, significant mapping density of the heterogametic sex-specific k-mers are still observed in the SDR. 3) Finally, more homogametic sex-specific *k*-mers are identified, which we observe in the 2.5✕ genome coverage subsets (Table 1) with the most dramatic increases observed in cannabis, hop, and brown bear. Encouragingly, the falsely identified homogametic sex-specific *k*-mers in the lower depth subsets do not map to the SDR, further showing the specificity of the SDR to the heterogametic sex and their ability to highlight sex-specific regions. Though nine species (Table S1) had less than the optimal low-depth sequencing ^18,94–96^ of 10✕ of the genome size, the severity of signal loss and *k*-mer noise varied, with the heterogametic sex-chromosome still identifiable with CBS-Discovery *k*-mers in those nine species (Figure 3, Figures S1-S2).

The beta regression model evaluating sex-specific *k*-mer mapping noise as a function of sequencing depth of coverage and number of isolates was only moderately predictive of noise (pseudo-R^2^ = 0.221). The main drivers of *k*-mer mapping noise, and therefore the ease of identifying the SDR with CBS-tools, are likely biological factors specific to each species, such as genetic diversity within the species, population structure amongst isolates used in CBS-Discovery, time of divergence between sex chromosomes, or conservation of SDR amongst heterogametic-sexed individuals. For CBS-Discovery to successfully return *k*-mers specific to the heterogametic-sex, the sample of homogametic isolates must capture the majority of genetic variation in the autosomes and PAR of the heterogametic-sex. Population structure within the species could suppress the signal if all males and all females are sampled from different populations. One different isolate within males or females will likely not have a huge impact, as all same-sex isolates will have to share the same genetic variants for them to appear as shared *k*-mers. The problem arises when none of the opposite sex shares that genetic variation for it to be filtered out as non-sex linked. The age of the sex-system could also impact sex-linked *k*-mer signals in two ways. If the sex-system is very-recently evolved, the sex chromosomes could still be similar to their ancestral autosomal state and potentially not have derived sufficient divergence between their homologous regions. If the SDR only recently stopped recombining, it might also be very small, which we consequently expect could lead to a small sex-specific list of *k*-mers.

These biological variables are difficult to quantify in a model, and are likely debated in some species, thus they have been left out of any predictive models. Sex-system is one of the few variables that can be accurately called for all 31 species used in this manuscript, however, the CBS-Discovery methodology should be agnostic to any difference in sex-system. When including sex-system as a categorical factor in the original beta regression model, we see no significant differences in *k*-mer noise in the XY systems (b = 0.307, z = 0.504, p = 0.614) or ZW systems (b=1.152, z = 1.663, p = 0.096) with UV set as the intercept (pseudo-R^2^ = 0.306). Furthermore, the estimated marginal means between the sex-systems found no significant differences in *k*-mer noise between pairwise comparisons of sex-systems (p > 0.1), supporting the sex-specific *k*-mer mapping results (Table 1, Table S3, Figure 3, Figure S1-S2).

### Considerations for other sex chromosome systems

CBS-tools handles variation in the number of sex-specific chromosomes in a species. As mentioned above, we recovered all Y chromosomes in the multiple systems of platypus and Japanese hop (Figure 3, Figures S1-S2). With UV systems, as both females and males contain a sex-specific chromosome, the heterogametic sex thus does not need to be identified via CBS-Discovery. However, CBS-tools can still provide a robust framework for identifying the U/V chromosomes in the genome assembly. For the two UV species sampled, we found the *k*-mer lists are nearly the same size (sex specific *k*-mer ratios: fire moss, 0.96; peat moss, 1.54). Any observed difference in the sex-specific *k*-mer ratio is likely to be driven by differences in the size of the U or V, or the sampling strategy. The U- and V-mers lists map to the expected chromosomes and scaffolds (Figure S1, Table S3), covering the entire U and V chromosomes in fire moss and the V in peat moss. We note that a scaffolded U chromosome assembly was not available for peat moss, however the peat moss U-mer list still maps to the U chromosome fragment available ^97^. Conversely, CBS-tools will struggle with XO or ZO systems, where sex determination is strictly dosage dependent ^98^ or if the previously Y-linked genes have moved to the X ^99^, as there are no regions or chromosomes specific to the heterogametic sex - just copy number variation.

Finally, species that are gynodioecious (female plants and bisexual plants), androdioecious (male individuals and bisexual individuals, though rare), or trioecious (unisex and bisexual individuals), could confound CBS-tools results ^100^. If a bisexual isolate is mis-phenotyped as a unisex male or female, all true sex-linked *k*-mers could be wiped out if the bisexual isolate shares the same or similar SDR as the heterogametic sex. In fact, one of the biggest challenges with CBS-Discovery is the hard requirement of known sex phenotypes of the isolates. Because the filtering process looks for *k*-mers shared by each sex that are not found in the other sex, any mislabeled individual will cancel out true sex-linked *k*-mers. For example, an XX female mistakenly labeled as a male in an XY system will cause Y-mers to be filtered out as autosomal noise since the female lacks a Y chromosome. Or in a ZW system, a female labeled as ZZ would cause CBS-Discovery to filter out the female-only *k*-mers as they can be detected in the “all male *k*-mers” database created with meryl union-sum ^59^ (Figure 1). Future versions of the CBS-tools pipeline may include pipeline updates that are more flexible with mis-phenotyped samples, however, for this version we recommend only using well vetted isolates.

### CBS-Phase tests for and guides phasing the sex-specific chromosome

Assemblers like HiFiasm ^84^, can occasionally produce a completely phased heterogametic assembly without using trio-binning, but without the parental information, the contigs end up as a technical artifact as chimeric across the haplotypes ^13,49^. In many cases, it is impossible to retrieve parental sequencing for trio-binning a long-read assembly, whether logistically or financially. While suitable for autosomes where the goal is not haplotype tracking in the progeny, chimerism still poses a problem for sex chromosomes. An SDR split across haplotypes can obscure sex-specific markers or alleles controlling sex determination, hindering research efforts. We have found the sex-specific *k*-mers from CBS-Discovery can be used to successfully identify Y or W related contigs in both haplotypes and manually phased when used alongside chromatin conformation sequencing and contact maps. The sex chromosomes in the reference genomes for cannabis and hop were previously joint-scaffolded and phased across haplotypes with mapped Y-mers in Carey et al. 2026. Here, the male-specific *k*-mers in both species obtained from CBS-Discovery all mapped overwhelmingly to their respective Y chromosome (Table 1) and the female-specific *k*-mers either do not map or map elsewhere in the genome. The same methods were also used previously in the Amborella reference with W-mers ^30^. With this previously phased assembly, we see the female-specific *k*-mers from CBS-Discovery that map to the genome map completely to the SDR (Table 1). No male-specific *k*-mers were returned.

For brown bear, WGS for both parental genotypes, “John” and “Oakley”, and Omni-C data, for their offspring “Adak”, were available, permitting the opportunity to directly compare CBS in two assemblies using different phasing strategies. Our a priori expectation is that the X and Y were pulled successfully into their respective haplotypes using the trio-binning approach (Armstrong et al., *in review*). Using CBS-Phase we correctly called the Y chromosome in the trio-binned assembly at both the scaffolded-scale (Table 1, Figure 3) and the contig-scale prior to the scaffolding step (Table S6), further demonstrating that the Y-mer list is a valuable test for phasing. For the Omni-C assembly, we mapped the Y-mers to identify Y-linked contigs. We mapped these Y-linked contigs with coverage >1x Y-mer coverage, as well as the contigs that represent the pseudoautosomal region (described below), to the trio-binned Y chromosome assembly and found 99.99% coverage. However, with Omni-C incorporation, the Y contigs demonstrably ended up split across the two haplotypes ^85^ (Table S7), and consequently needed to be manually moved into one haplotype prior to final scaffolding. With the additional joint-scaffolding approach, we were able to vet the Y-mer hinted contigs within the Hi-C contact map, identify the correct piece of the PAR for the Y, as well as identify the X chromosome contigs (Figure S3, Table S8). Like the Y chromosome, contigs for the X were also split between the haplotypes in the contig assembly (Table S8). Using the combination of CBS-Phase and joint-scaffolding, we recovered 99.97% of the X chromosome and 98.07% of the Y in the Omni-C assembly, relative to the trio-binned. There may be other approaches suitable for identifying the X-linked contigs, such as through alignments to a closely-related reference. Importantly, many unexplored sex chromosomes are likely to lack a closely-related neighboring species, or syntenic relationships may quickly dissolve, making the CBS-Phase and joint-scaffolding a pragmatic approach.

Future versions of CBS-tools may attempt to automate phasing sex chromosomes across haplotypes in non trio-binned assemblies. However, as genome sequencing and assembly technologies continue to improve, testing for phasing may become a less necessary step. Indeed, assemblies are falling out as continuously assembled from telomere-to-telomere ^15,16,101^. Currently, the input to generate T2T assemblies may be cost-prohibitive, and with the wealth of data being generated, at least for the foreseeable future, CBS-Phase will be of benefit to those still manually phasing sex chromosome contigs. CBS-Phase can also help to vet the sex-specific chromosome in a genome assembly. This is of benefit because there are currently few QC metrics available specifically for the sex chromosomes. Indeed, even the standard of BUSCO for vetting an assembly lacks genes on the sex chromosomes to help guide the quality, at least in mammals ^102^.

### CBS-Frontier estimates SDR boundaries and visualizes k-mer mapped genes

The final frontier before engaging in analyses of the sex chromosomes is delimiting the boundary between the PAR and the SDR. This boundary can be a difficult location to identify, even in the most well-studied species, humans ^103^. PARs can also be variable within a species ^104^. We used CBS-Frontier to estimate SDR boundaries for nine species representing different sex-system and sex-chromosome characteristics across plants and animals and visualize *k*-mer linked genes, some of which are known sex determination candidates. The changepoint analysis reliably called the PAR/SDR boundaries consistently for species with the previously reported SDRs (Table S1, Table S5). With the exception of white mulberry, the overwhelming majority of *k*-mers from the heterogametic sex fall within these boundaries on the sex-specific chromosomes (Table 1, Table S5, Figure 3). The SDR boundaries were not outlined previously in the threespine stickleback assembly ^70^. However, within the boundary called by CBS-Frontier, we found the reported sex determination gene *amhy* (Figure 5). In brown bear, CBS-Frontier called the PAR/SDR boundary at 24.23 Mb, making *SHROOM2* (Figures 3-5) the first gene in the PAR, which has been previously demonstrated in other Carnivora species ^66^. Within this boundary, we find Y-mers mapped to the known sex-linked gene *SRY* ^9,10^. The Salicaceae family is arguably the most notable clade to test the efficacy of CBS-tools because of their complex evolutionary background. Poplars (*Populus* sp.: black cottonwood, balsam poplar, white poplar) and willows (*Salix* sp.) contain mostly dioecious species that may have a shared origin, yet they have undergone several translocations of the SDRs, jumping within and between chromosomes, and in some cases switching from XY to ZW ^105–107^. These translocations involve the functionally validated sex-determining gene *ARR17* ^92,108^ In black cottonwood the SDR region contains 5 annotated gene models ^75^ and between 16,290,271-16,307,190 bp, there are 5 copies of *ARR17* partial repeats with varying exons from the original *ARR17* model. One gene previously suggested to be in the SDR, a hypothetical protein (PtStettler14.18G127500; 16,214,199-16,215,600bp), was found marginally outside of the boundary called by CBS-Frontier (16,215,000bp; Table S5). However, this gene had no synonymous substitutions per synonymous site (Ks of 0) when comparing the X:Y orthologs in Zhou et al. (2020), suggesting there is no divergence between the X and Y copies. Increased Ks is a typical observation in sex-linked genes ^109^, and the lack of divergence, and our CBS boundary call, suggests this gene may not be sex-linked. Similarly, we found a smaller SDR in *S. dunni* than was reported. He et al. (2021) called the PAR/SDR boundary using *F_ST_*to estimate genetic differentiation between sexes ^54^. High *F_ST_* between the sexes is a signature of sex-linkage, but it can also be due to the strong selection at the PAR boundary ^29^. Alternatively, the SDR boundaries for these species could differ across genotypes and those used in CBS have the smaller SDR size. Users of CBS-tools are as such cautioned to holistically examine these results within the context of their focal species and other subsequent analyses.

**Figure 5.**
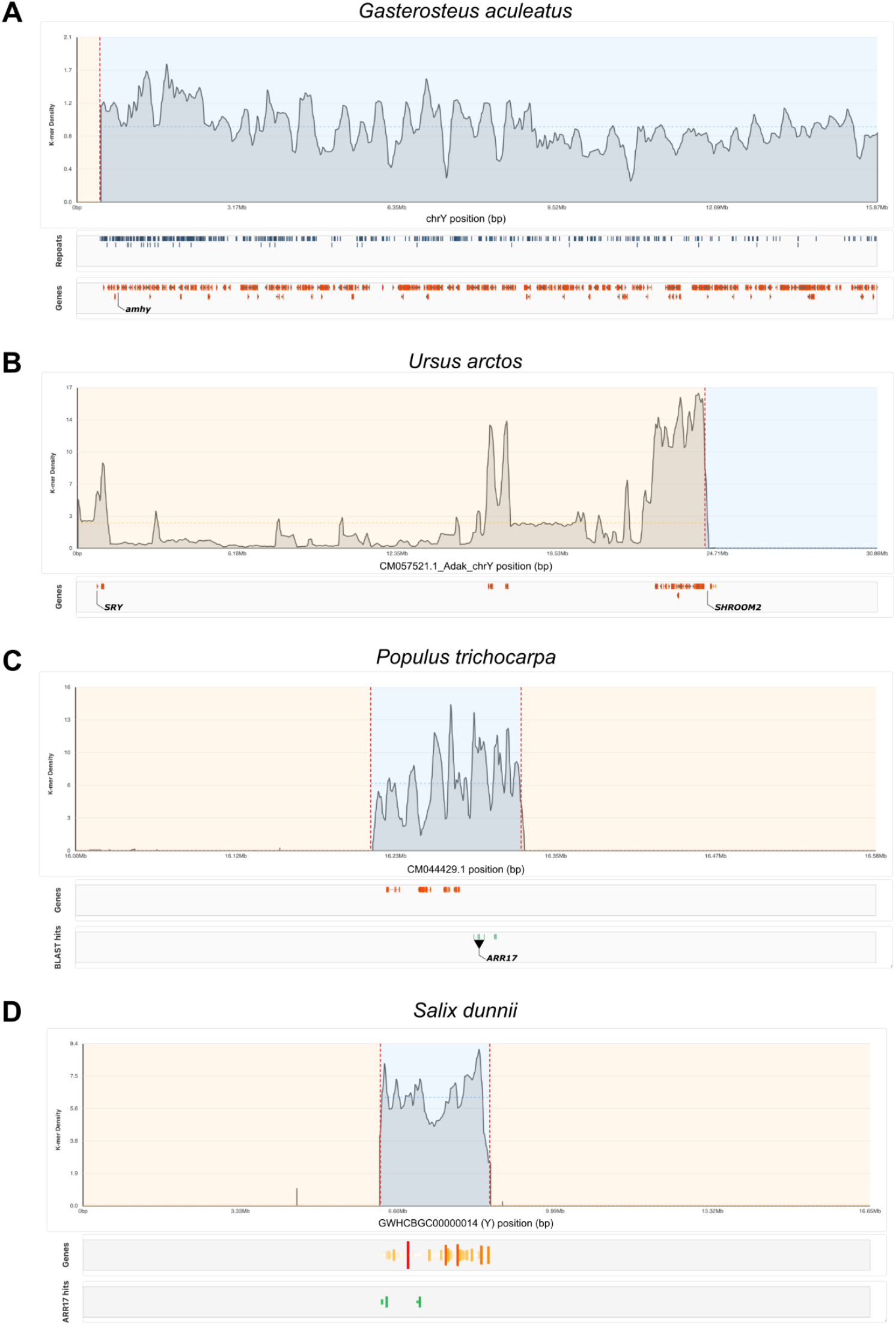
Sex-specific *k*-mers in four species mapped to genes located within the sex-determining region (SDR) in CBS-Frontier. A) *Amhy* in *Gasterosteus aculeatus* (threespine stickleback). B) *SRY* in *Ursus arctos* (brown bear). The SHROOM2-like gene (LOC125282542) location is also marked at the start of the PAR boundary. C) five partial hits to *ARR17* in *Populus trichocarpa* (black cottonwood), and D) six partial hits to *ARR17* in *Salix dunnii* (willow). *K*-mer peaks are plotted at 50kb resolutions and gene models shown are those linked to sex-specific *k*-mers above a 50 *k*-mer threshold. Boundaries estimated with changepoint analyses (Table S5) are denoted with vertical dotted lines in each panel.

## Conclusions

The evolution of sex chromosomes is a phenomenon that spans taxonomic kingdoms. Here we developed a *k*-mer based toolkit that works to examine sex chromosomes across the broadest diversity of pairs. The classic approach for identifying sex chromosomes is through cytology. However, many sex chromosome pairs are homomorphic at that scale, and cannot be reliably determined ^5,6^, in addition to the other challenges associated with cytological approaches (e.g., needing root tips or cell cultures; large chromosome numbers with variable sized chromosomes). Some computational pipelines to identify the sex chromosomes are also available ^8^. Analyses that examine patterns of allelic segregation are powerful for identifying sex-linked sequences, but can rely on a pedigree that includes a cross of parents with sufficient genetic diversity, and subsequent phenotyping of the offspring ^110^. One challenge is that reproduction and sex expression can take a long time, or be simply impossible, in some species. Many other pipelines that have been developed to examine sex chromosomes typically rely on having an existing genome reference to calculate differences in coverage between the sexes or run population genomic analyses ^111–114^. Our data clearly show that standard approaches to genome assembly will result in improper phasing and/or the misassembly of the sex chromosomes, which complicates the interpretation of these analyses, especially with regards to studying the SDR. More recently, software has relied on the genome assembly itself to identify the sex chromosomes ^115^. However, this approach relies on the heterogametic sex being known and is currently only recommended for use in vertebrates ^115^. Moreover, our analyses emphasize that using small numbers of isolates can be problematic for identifying the sex chromosomes (Table 2, Figure 4). *K*-mer based tools have shown great potential for a variety of genomic issues ^60^, including examining sex chromosomes (e.g., ^30,39,115–118^). A major advantage of the CBS-tools approach is that the same *k*-mer list can address many aspects of sex chromosome biology, broadly overcomes limitations present in other sex chromosome analysis approaches, and is largely universal across the tree of life. Despite hundreds to thousands of independent origins across the tree of life, sex chromosomes exhibit similar structural characteristics that can be successfully explored with *k*-mer based analyses.

## Supplemental

**Figure S1.** Sex-specific *k*-mers mapped to 22 dioecious plant species reference genomes.

**Figure S2.** Sex-specific *k*-mers mapped to 9 animal species reference genomes.

**Figure S3.** Hi-C contact map to identify the X- and Y-linked contigs for *Ursus arctos* “Adak”.

**Table S1.** Summary of species and their sex chromosome systems tested with CBS-tools. Species with closely related, but non-conspecific, references have been marked with NCS under Reference Assembly.

**Table S2.** Isolates used in CBS-Discovery from WGS accessions in Table S1.

**Table S3.** Sex-specific *k*-mers mapping rates to the remaining species not selected as examples in Table S1.

**Table S4.** *A. trichopoda* and *G. aculeatus* sex-specific *k*-mer counts from all replications when subsetting the number of isolates used in CBS-Discovery.

**Table S5.** SDR boundary values determined by changepoint analysis in the seven representative species in Figure 3.

**Table S6.** Y-mer mapping on the trio-binned assembly for *Ursus arctos* “Adak”.

**Table S7.** Y-mer mapping on the Omni-C incorporated assembly for *Ursus arctos* “Adak”.

**Table S8.** Contigs identified for the XY chromosomes in the Omni-C incorporated assembly for *Ursus arctos* “Adak” using the joint-scaffolding approach.

## Data availability

All data used in this manuscript were sourced from existing, publicly-available, resources. The assembly and whole-genome sequencing accessions can be found in Table S1 and Table S2. Additional results in support of this manuscript, including the final sex-specific k-mer lists, can be found on Zenodo (DOI: 10.5281/zenodo.22836531).

## Code availability

CBS-tools is available on GitHub at https://github.com/HudsonAlpha/CBS-tools and the interactive *k*-mer plotting tool, CBS-Frontier, is available at https://github.com/HudsonAlpha/CBS-Frontier.

## Acknowledgements

We would like to thank Nicole Stark for testing early versions of this methodology. We would also like to thank the HudsonAlpha IT Department for their research computing support.

Funding support for this work was provided by the National Science Foundation (NSF) Graduate Research Fellowship Program (L.M.A.), the NSF IOS-PGRP CAREER no. 2239530 (A.H.), the NSF EDGE no. 2421469 (A.H.), and the U.S. Department of Agriculture National Institute of Food and Agriculture Postdoctoral Fellowship (USDA NIFA) no. 2022-67012-38987 (S.B.C.).

## Author Contributions

L.W.: Data curation, Formal analysis, Software, Visualization, Methodology, Writing-original draft, Writing-review & editing

L.M.A.: Methodology, Software, Visualization, Writing-original draft, Writing-review & editing

P.C.B.: Resources, Writing-review & editing

E.E.A.: Resources, Writing-review & editing

A.H.: Conceptualization, Resources, Funding acquisition, Writing-review & editing

S.B.C.: Conceptualization, Data curation, Formal analysis, Visualization, Methodology, Funding acquisition, Resources, Writing-original draft, Writing-review & editing

