## Supplemental Figures 1-3 for "Identifying, phasing, and structurally annotating sex chromosomes for genome assemblies using CBS-tools"

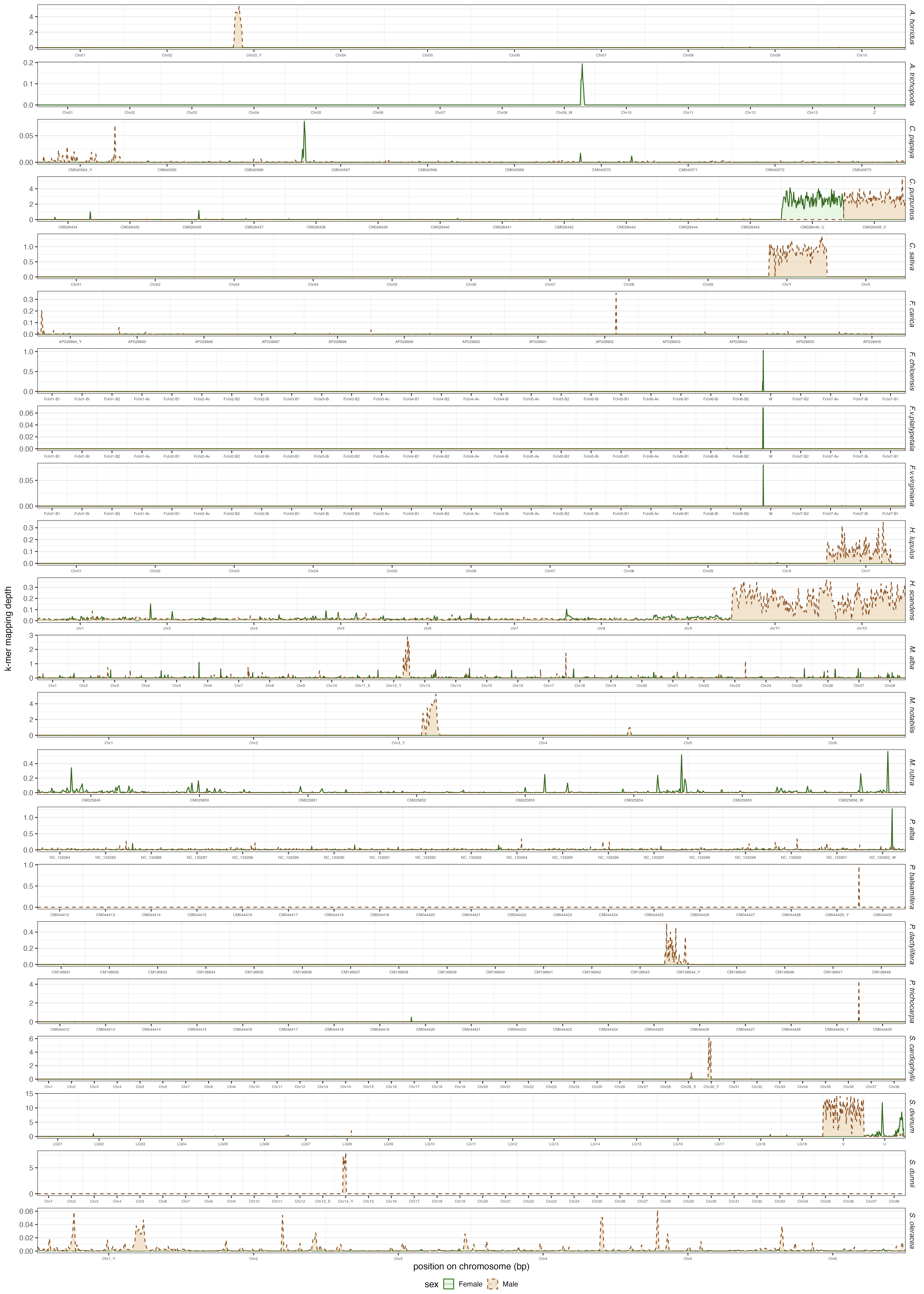

**Figure S1. Mapping sex-specific k-mers to reference genomes in 22 plant species (Table 1).** Only chromosomes are shown, each binned into 100 equal sized windows for averaging k-mer mapping depth within bins. Mapping depth is plotted as averages over base pairs within bins. For haplotype resolved assemblies, only one haplotype is shown but sex chromosomes from both are shown.

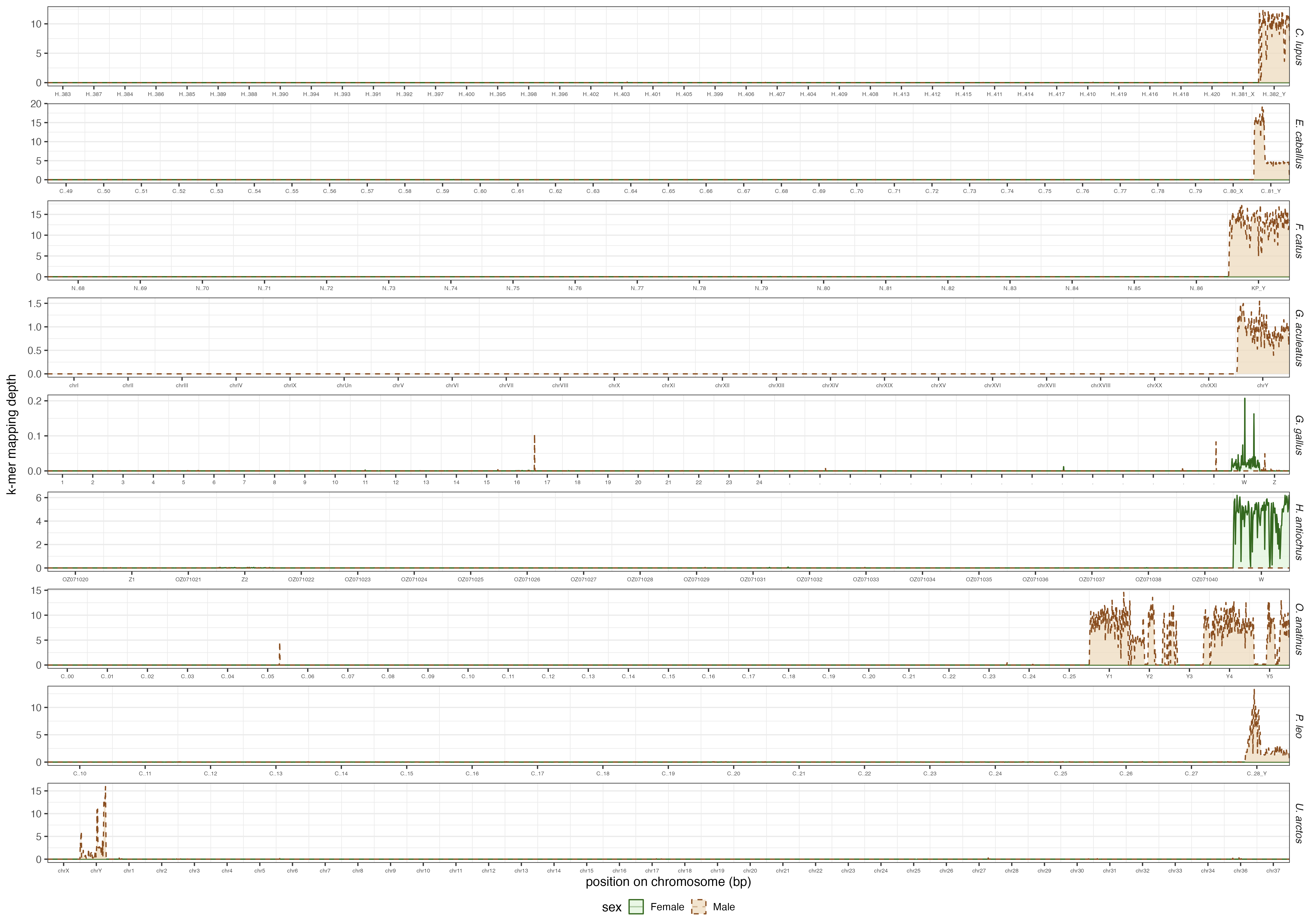

**Figure S2. Mapping sex-specific k-mers to reference genomes of 9 animal species (Table 1).** Only chromosomes are shown, each binned into 100 equal sized windows for averaging k-mer mapping depth within bins. Mapping depth is plotted as averages over base pairs within bins.

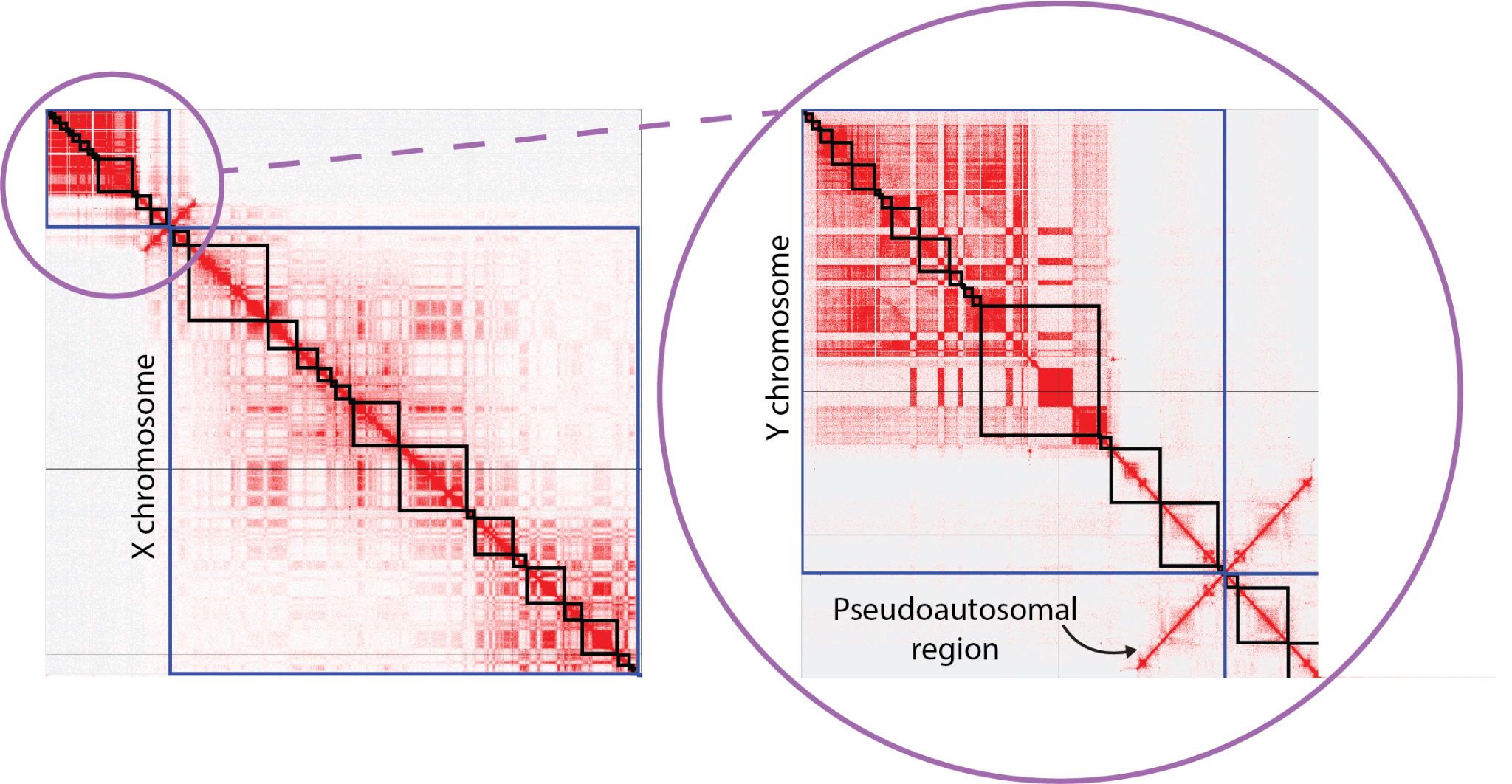
**Figure S3. Hi-C contact map to identify the X- and Y-linked contigs for *Ursus arctos* "Adak".** We used the joint-scaffolding approach (described in the Methods) to validate the contiguity of the Y-mer mapped contigs and identify the correct piece of the pseudoautosomal region (PAR) for the Y (Table S7), as well as to identify the X-linked contigs (Table S8). This assembly used Omni-C data to phase the contigs, rather than trio-binning and we identified contigs for the X and the Y chromosomes in both haplotypes of the assembly (Table S8). The PAR is apparent through the offdiagonals in the Hi-C contact map. The black boxes represent individual contigs, while the blue box denotes the boundary of the scaffolded region (ideally as a chromosome or pseudomolecule representation of one).
